# Feature-Driven Whole-Tissue Imaging with Subcellular Resolution

**DOI:** 10.1101/2025.01.23.634600

**Authors:** Jinlong Lin, Zach Marin, Xiaoding Wang, Hazel M. Borges, Pierre-Emmanuel Y. N’Guetta, Xuemei Luo, Baylee A. Porter, Yuanyuan Xue, Md Torikul Islam, Tai Ngo, Arin B. Aurora, Hu Zhao, Suzanne D. Conzen, Sean J. Morrison, Shuang Liang, Zhenyu Zhong, Lori L. O’Brien, Kevin M. Dean

## Abstract

Understanding how subcellular structures and events shape tissue-wide phenomena remains a central challenge in biology. We present Multiscale Cleared Tissue Axially Swept Light-Sheet Microscopy (MCT-ASLM), a platform combining cm-scale imaging with targeted high-resolution interrogation of intact tissues in human-guided or autonomous modes. Capable of capturing fields of view up to 21 mm at micron-scale resolution, MCT-ASLM can seamlessly transition to submicron (~300 nm) isotropic resolution for targeted imaging. This versatility enables comprehensive studies of hierarchical organization and spatially complex processes, including mapping neuronal circuits in rat brains, visualizing glomerular innervation in mouse kidneys, and examining metastatic tumor microenvironments. By bridging subcellular to tissue-level scales, MCT-ASLM offers a powerful method for unraveling how local events contribute to global biological phenomena.

## Introduction

Tissue development, structure, and function manifests from the complex organization of diverse cell types acting in unison^1^. Yet, studying cell biological phenomena in both developing and terminally differentiated tissue contexts is challenging because cells are influenced by local environmental cues that vary significantly throughout a tissue^2^. These cues, including cell-cell interactions, extracellular matrix composition, cytokines, hormones, and more, create a spatially varying microenvironment that shapes cellular behavior and responses. For example, both sympathetic and sensory nerves modulate kidney function through precise neural control of blood flow, filtration, and electrolyte balance. However, kidney innervation during development exhibits significant spatial heterogeneity, with differences in nerve density and patterning across regions^3^. Understanding how such intricate spatial relationships emerge and influence biological function requires quantitative imaging and analytical tools capable of linking molecular and cellular events to broader tissue-wide structures and functions across vast spatial scales.

Current methods for volumetrically imaging large biological specimens often struggle to meet the dual demands of resolving fine subcellular structures while capturing larger tissue architectures^4^. Light-sheet fluorescence microscopes (LSFM), for example, typically operate at a single magnification, achieving lateral resolutions of ~300 nm^5^. However, obtaining a large field of view (FOV) in LSFM often compromises axial resolution, which is particularly problematic in biological contexts where tissue architectures, and the cells that constitute them, extend in diverse and complex spatial directions^3^. This reduction in axial resolution obscures critical three-dimensional details necessary for quantitatively evaluating cellular behavior and its role in broader biological outcomes^6^. An ideal microscope should operate seamlessly across multiple spatial scales, offering a large FOV and isotropic resolution to faithfully capture the intricate three-dimensional organization of tissue architecture and cellular morphology within complex biological systems.

Recent advancements in LSFM have introduced the capacity to operate across multiple spatial scales. For example, the Hybrid Open-Top Light-Sheet Microscope utilized orthogonal and non-orthogonal imaging paths to achieve multiscale imaging across an expansive 10 × 75 × 120 mm volume^7^. However, its high-resolution non-orthogonal imaging module suffered from an axial resolution of 2.91 microns, which is 6.3 times worse than its lateral resolution of 0.45 microns. A variant of Axially Swept Light-Sheet Microscopy (ASLM) was designed with a detection path equipped with a motorized flip mirror to enable imaging at different magnifications^8^. This system allowed researchers to image entire organoids and evaluate immune-cancer cell interactions in zebrafish xenografts. However, it was limited to only two scales, and its imaging volume of 1.3 × 1.3 mm was relatively small compared to the size of typical mouse and human tissues. Furthermore, despite leveraging the ASLM acquisition format, a significant disparity remained between its lateral (390 nm) and axial (650 nm) resolutions. In parallel, a vibratome-equipped multiscale light-sheet fluorescence microscopy system was developed that could flexibly adjust its illumination and detection optics, with magnifications varying from 1.26x to 12.6x. Also equipped with an ASLM detection scheme, this system achieved a resolution of 0.95 μm × 0.95 μm × 2.1 μm along the X-Y-Z axes^9^. While these advancements offer notable improvements, they remain insufficient to address how larger tissue microenvironments influence biological function at sub-cellular scales.

Another significant challenge associated with high-resolution tissue imaging is the time required to acquire data and the sheer scale of the resulting datasets, much of which may not directly pertain to the biological question of interest^10,11^. These issues become particularly pronounced when attempting to identify rare features, such as metastases, or to comprehensively evaluate biological events in a statistically robust and informative format. Smart microscopy addresses these challenges by integrating automated image analysis with adaptive acquisition routines, allowing imaging to be dynamically guided by the biological content of the specimen^1,12^. By leveraging real-time data analysis, smart microscopy focuses imaging resources only on regions of interest, reducing both acquisition time and data overhead. Here, we sought to integrate the strengths of multiscale imaging with a versatile acquisition framework that supports both human-in-the-loop and autonomous imaging modes. Using this approach, large-scale tissue volumes are first imaged at low magnification to provide a global overview, enabling candidate features to be identified on-the-fly with classical computer vision techniques that do not require training or fine-tuning convolutional neural network-based models. These features are then automatically interrogated at high resolution, achieving ~300 nm isotropic resolution. This capability is realized through a 4-axis light-sheet microscope that combines macroscale and nanoscale imaging modules, shared sample positioning, and completely reconfigurable software^13^. We demonstrate the generality of this approach by imaging renal, genitourinary, hepatic, neural, pulmonary, and skeletal tissues. Altogether, our approach enables comprehensive, high-replicate evaluation of biological features, greatly accelerating the pace of discovery across a broad range of applications.

## Results

### Imaging Across Scales with MCT-ASLM

To address the challenges of multiscale imaging, we developed Multiscale Cleared Tissue ASLM (MCT-ASLM), a versatile LSFM platform designed for comprehensive tissue imaging (**Figure 1A, Figure S1, Supplementary Tables 1 and 2)**. MCT-ASLM features a 4-axis geometry that integrates both macroscale and nanoscale imaging modules, with compatibility across a wide range of optical clearing solvents (1.333 to 1.56). Both modules leverage the ASLM acquisition format to achieve high-resolution imaging across a large FOV^14^. The macroscale module offers a continuously variable magnification of 0.63x to 6.3x, enabling the acquisition of FOVs ranging from 21.1 mm x 21.1 mm to 2.1 mm x 2.1 mm, respectively, when paired with standard 15 mm cameras^15^. At 6x magnification, the macroscale module achieves a resolution of 2.4 μm laterally and 4.2 μm axially in aqueous environments before deconvolution (**Figure S2**), which improves to ~2 μm laterally and 3.2 μm axially after deconvolution (**Figure S3, Supplementary Table 3**)^16^. These resolutions are preserved in high-refractive index solvents, as measured on CF555-antibody aggregates in BABB-cleared lung specimens (**Supplementary Tables 4, 5, 6, and 7**). While sub-Nyquist lateral spatial sampling is not suitable for resolving diffraction-limited objects, magnifications below 6x remain effective for detecting larger features, such as single cells and populations of cells. The nanoscale module of MCT-ASLM provides a magnification range of 32x to 38x, depending on the refractive index of the immersion medium. In aqueous environments, this module achieves a FOV of 468 µm × 468 μm and an isotropic resolution of 470 nm (**Figure S4**). After deconvolution, the average lateral and axial resolution improves to 380 and 333 nm (**Figures S5**), respectively. This level of resolution is particularly advantageous for investigating sub-cellular structures and morphological features of single cells within their native tissue environments. While deconvolution, when judiciously implemented, can enhance both resolution and contrast, all data presented in this manuscript is shown in its raw, unadulterated format.

**Figure 1.**
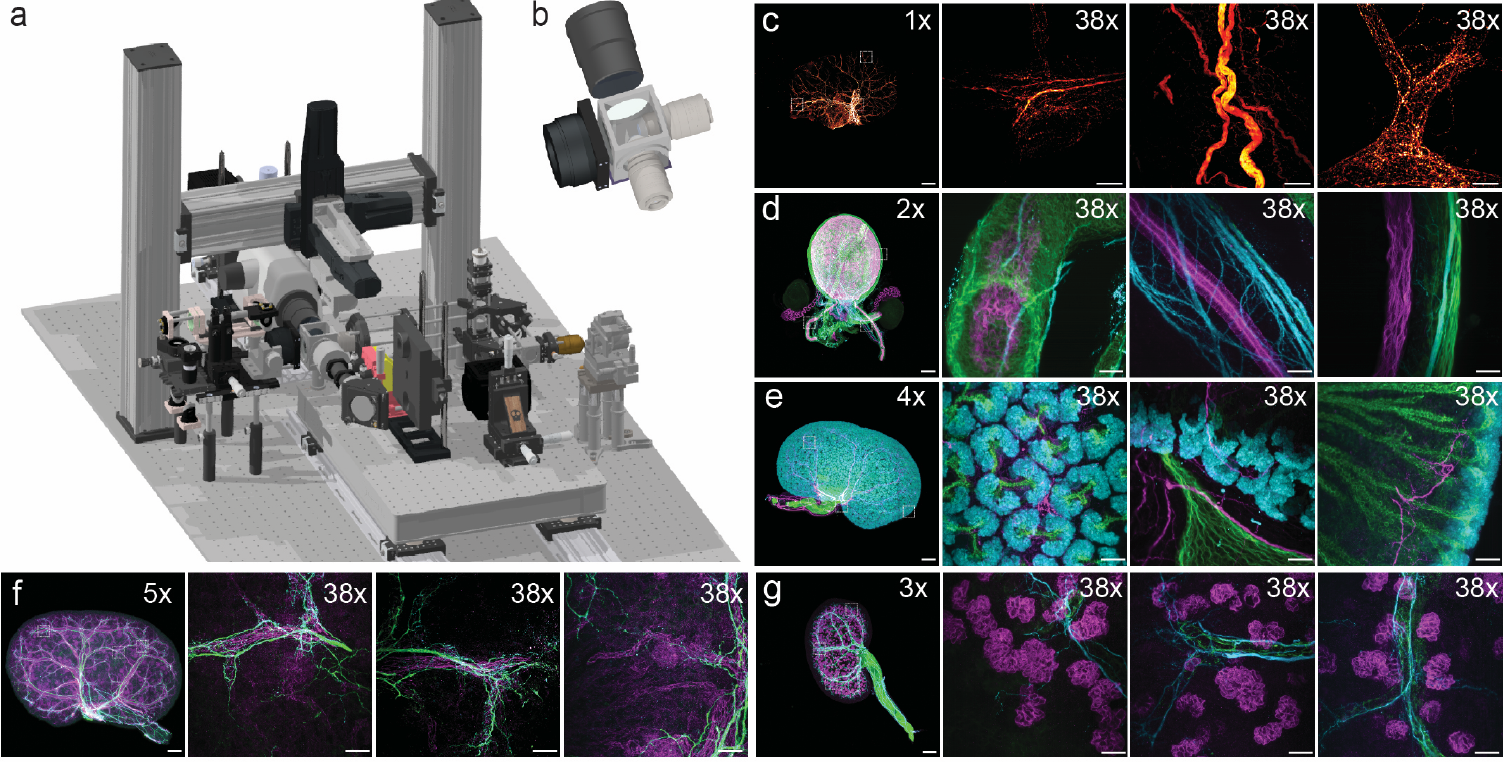
Microscope Design and Multiscale Biological Imaging. **a)** CAD rendering of the MCT-ASLM system. **b)** Close-up view of the sample chamber and illumination axes. **c)** Mouse liver lobe stained for tyrosine hydroxylase, imaged at 1x and 38x magnifications. High magnification reveals detailed structures within the liver’s autonomic nervous system. **d)** Mouse bladder stained for SMA (green), TUBB3 (cyan), and cytokeratin (magenta), imaged at 2x and 38x magnifications. Smooth muscle fibers and neuronal tracts are visible within the bladder wall and urethra. **e)** Mouse kidney stained for cytokeratin (green), SIX2 (cyan), and TUBB3 (magenta), imaged at 4x and 38x magnifications. SIX2 labels nephron progenitors, while TUBB3 highlights neuronal components. **f)** Mouse kidney stained for TUBB3 (green), Synapsin (cyan), and CD31 (magenta), imaged at 5x and 38x magnifications. High-resolution imaging reveals interactions between neuronal synaptic structures and vasculature. **g)** Mouse kidney stained for SMA (green), TUBB3 (cyan), and Podocin (magenta), imaged at 3x and 38x magnifications. Subcellular details, including podocyte networks, are evident. Scale bars: C, 1x = 1000 µm, 38x = 50 µm; D, 2x = 500 µm, 38x = 50 µm; E, 4x = 250 µm, 38x = 50 µm; F, 5x = 200 µm, 38x = 50 µm; G, 3x = 333 µm, 38x = 50 µm. See methods for additional details on labeling and clearing methods.

Each imaging module is equipped with independent focusing stages but share a common 4D (XYZθ) sample positioning system. The sample is positioned from above within a large chamber featuring optical windows for the macroscale module and silicone gaskets on the remaining sides to allow mechanical access for two multi-immersion dipping objectives (**Figure 1B**). With the working distance of the optics, travel ranges of the stages, and specimen chamber, MCT-ASLM can image specimens as large as 20 mm × 20 mm × 30 mm, so long as they are sufficiently optically clear—a 13-fold larger volume than the Hybrid Open-Top Light-Sheet Microscope. The system is controlled using navigate, a Python-based software package that enables seamless switching between the macroscale and nanoscale modules (**Supplementary Notes 1 and 2**). This allows volumetric interrogation of biological specimens, such as an intact mouse femur (**Figure S6**) or a mouse lung with spontaneously formed metastatic melanoma colonies^17^ (**Figure S7**), at magnifications ranging from 0.63x to 38x. With navigate, users can switch effortlessly between different image acquisition modes, leveraging both human-in-the-loop control for guided imaging and autonomous formats for efficient high-throughput data acquisition.

### Human-Guided Imaging of Hepatic and Urogenital Systems with MCT-ASLM

To demonstrate the versatility and power of MCT-ASLM, we applied the system to image diverse biological specimens in a human-driven acquisition format. In **Figure 1c**, we imaged a BABB-cleared liver stained for tyrosine hydroxylase, highlighting the autonomic nerves associated with the hepatic portal vein, a critical component of the hepatic nervous plexus. This dataset, acquired over a 10 × 6 ×3 mm liver lobule, revealed axonal protrusions branching throughout the hepatic tree, where the macroscale mode was used to identify regions of interest. These areas were further interrogated with the nanoscale module to uncover bulbous axonal features and dense axonal tracts. **Figure 1d** showcases imaging of a male mouse bladder and gonads stained for smooth muscle actin (SMA, green), TUBB3 (cyan), and cytokeratin (magenta), highlighting the intricate organization of smooth muscle fibers and nerves traversing the bladder walls and urethral tube. This structural detail highlights the intricate interplay between neuronal and muscular components in urogenital tissues, where SMA-rich layers form the structural and contractile framework, while neuronal projections traced by TUBB3 play critical roles in regulating coordinated physiological processes, including bladder contraction and relaxation during micturition. In **Figures 1e, 1f, and 1g**, we focused on mouse kidneys, leveraging the macroscale and nanoscale modes to resolve diverse cellular and structural features. In **Figure 1e**, imaging at 4x and 38x magnifications revealed cytokeratin (green) labeling of the collecting duct system, SIX2 (cyan) marking nephron progenitors, and TUBB3 (magenta) highlighting neuronal axons which project to the cortical progenitor niches and traverse areas of nephron differentiation. **Figure 1f** emphasizes the interplay between TUBB3 (green), synapsin (cyan), and CD31 (magenta), with axons and synaptic features associated with glomeruli and vasculature. Finally, **Figure 1G** demonstrates the labeling of SMA (green), TUBB3 (cyan), and Podocin (magenta), where high magnification unveiled detailed interactions between vasculature, nerves, kidney filtration.

### Multiscale Evaluation of Axonal and Neuronal Features in Rat Brains

The rat brain has become an increasingly important model in neuroscience, offering closer parallels to human brain structure and function than the widely used mouse model. With a volume of approximately 1750 mm^3^, the rat brain is over four times larger than the mouse brain, which measures ~400 mm^3^, posing significant challenges for imaging. Capturing large-scale structural features with synapse-scale detail requires advanced tools with large working distances and fast imaging mechanisms to efficiently span vast spatial scales without compromising resolution. **Figure 2a** illustrates a stereotaxic AAV-injected 12-week-old rat brain cleared with TESOS; axonal tracks spanning more than 13 mm in length are readily observed, while high-resolution imaging resolves synaptic-like features. In a complementary experiment, we imaged a whole rat brain with sparsely labeled neurons, a task made particularly challenging by the brain’s large size and the sparse distribution of the labeled cells (**Figure 2b**). To address this, the entire brain was first imaged at 1x magnification, providing a global overview. From this dataset, a region of interest was identified and further interrogated at 6x magnification. Within this region, multiple positions were subsequently selected for fully automatic multi-position imaging with the nanoscale module, where the microscope updated imaging parameters and acquired high-resolution data of the selected neurons.

**Figure 2.**
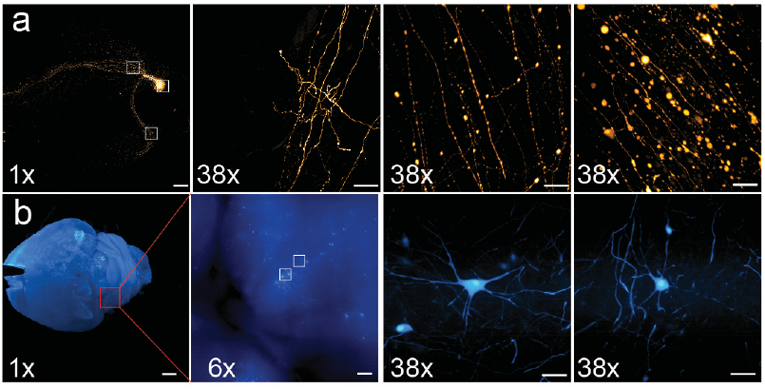
Multiscale imaging of neuronal architectures in the rat brain. **a)** Rat brain expressing an mScarlet reporter, imaged at 1x and 38x magnifications. Long-range axonal projections spanning over 13 mm are visualized, and synaptic-like features are resolved at high magnification. **b)** Imaging sparse neurons in a rat brain. The entire brain was imaged at 1x magnification to provide a global overview, followed by 6x magnification for region-specific interrogation and fully automatic multi-position imaging at 38x magnification to capture high-resolution neuronal details. Scale bars: A, 1x = 1000 µm, 38x = 50 µm; B, 1x = 1000 µm, 6x = 170 µm, 38x = 50 µm.

### Mapping Metastatic Colonization with MCT-ASLM

Another area where multiscale imaging proves particularly advantageous is in the study of metastatic colonization. Metastasis is an inefficient process during which cancer cells leave the primary tumor, enter the vasculature, extravasate into distant tissues (e.g. bone), adapt to a new microenvironment, and and glomeruli essential for proliferate to form metastatic lesions^18-20^. Thus, locating and further interrogating distant metastases in whole tissues isolated from in vivo cancer models requires the ability to capture both the broad spatial distribution of metastatic cells and their detailed interactions with the surrounding tissue microenvironment. In **Figure 3**, we evaluated infiltrating lobular breast cancer metastases originating in the mammary gland and later found in a mouse femur. In this model, the mouse bone was stained for laminin (green) to highlight vasculature and luciferase (magenta) to label metastatic cells, and imaged at 1x, 6x, and 38x magnifications. At lower magnifications, the metastatic burden and its distribution in the femur (1x, **Figure 3a**) and across the bone microenvironment (6x, **Figure 3b**) were evident, while 38x magnification (**Figure 3c, d**) provided insight into the spatial relationship between individual metastatic cells (magenta) and the surrounding microvasculature (green, **Video S1**). This level of detail could be used to gain insight into potential drivers of metastatic colonization, such as laminin-rich vascular niches, underscoring the capability of MCT-ASLM to interrogate both widespread tissue organization and intricate cellular features within complex biological systems.

**Figure 3.**
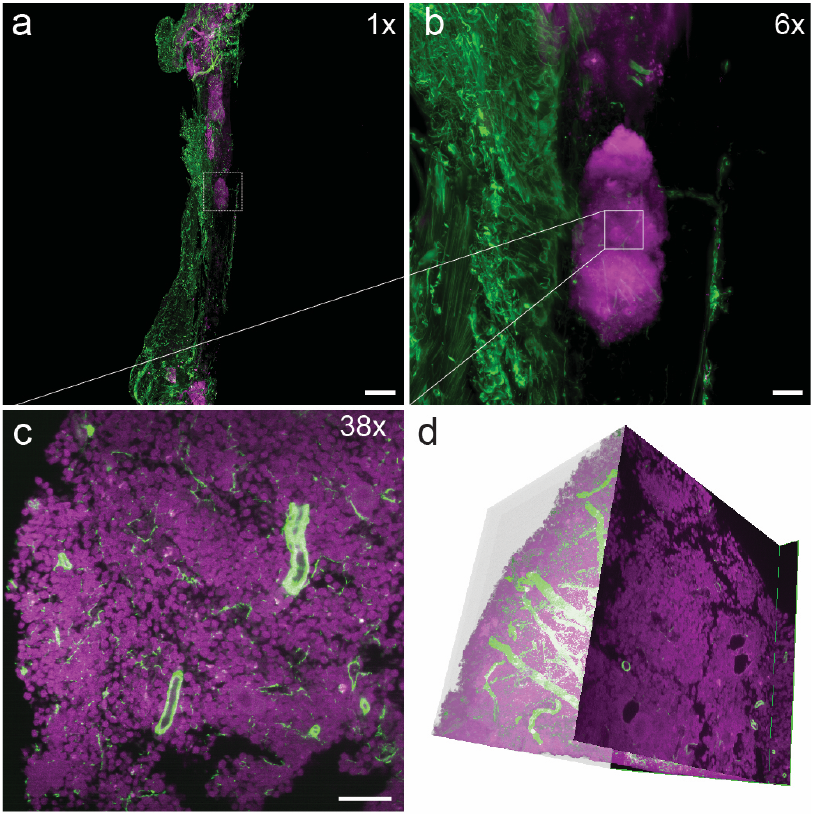
Imaging metastatic colonization with MCT-ASLM. Mouse bone stained for laminin (green) and luciferase (magenta), imaged at **a)** 1x, **b)** 6x, and **c)** 38x magnifications. **d)** A 3D rendering of the 38x magnification reveals a dense metastatic tumor mass and its proximity to the surrounding vasculature. Scale bars: 1x = 1000 µm, 6x = 170 µm, 38x = 50 µm.

### Autonomous Multiscale Imaging with MCT-ASLM

Next, we sought to operate MCT-ASLM in a smart microscopy format^1,11,12^. Here, we leveraged navigate’s feature framework, which enables the creation of custom protocols that can be arbitrarily chained together into reusable and image-guided acquisition routines (**Supplementary Note 3**)^13^. For instance, in **Figure 4A**, we perform automatic multi-resolution tiling of a lung from Pf4-cre; Ai47^loxpStop^ mice stained with antibodies against GFP, which reflects the PF4/CXCL4 expression pattern in platelets and megakaryocytes, and endomucin, a cell surface endothelial cell marker. CXCL4 is thought to be a driver of platelet-endothelial interactions that modulates pulmonary homeostasis and inflammation. To the best of our knowledge, this represents the first demonstration of automated tiling between independent imaging paths. Using navigate, MCT-ASLM automatically detected tissue boundaries at 6× magnification with the macroscale module, generated a sequence of positions to image at 38× magnification with the nanoscale module, switched operating modes, and acquired data in a tiled format. Such smart tiling formats streamline workflows and minimize data overhead by focusing imaging on regions of interest, avoiding the acquisition of empty space.

**Figure 4.**
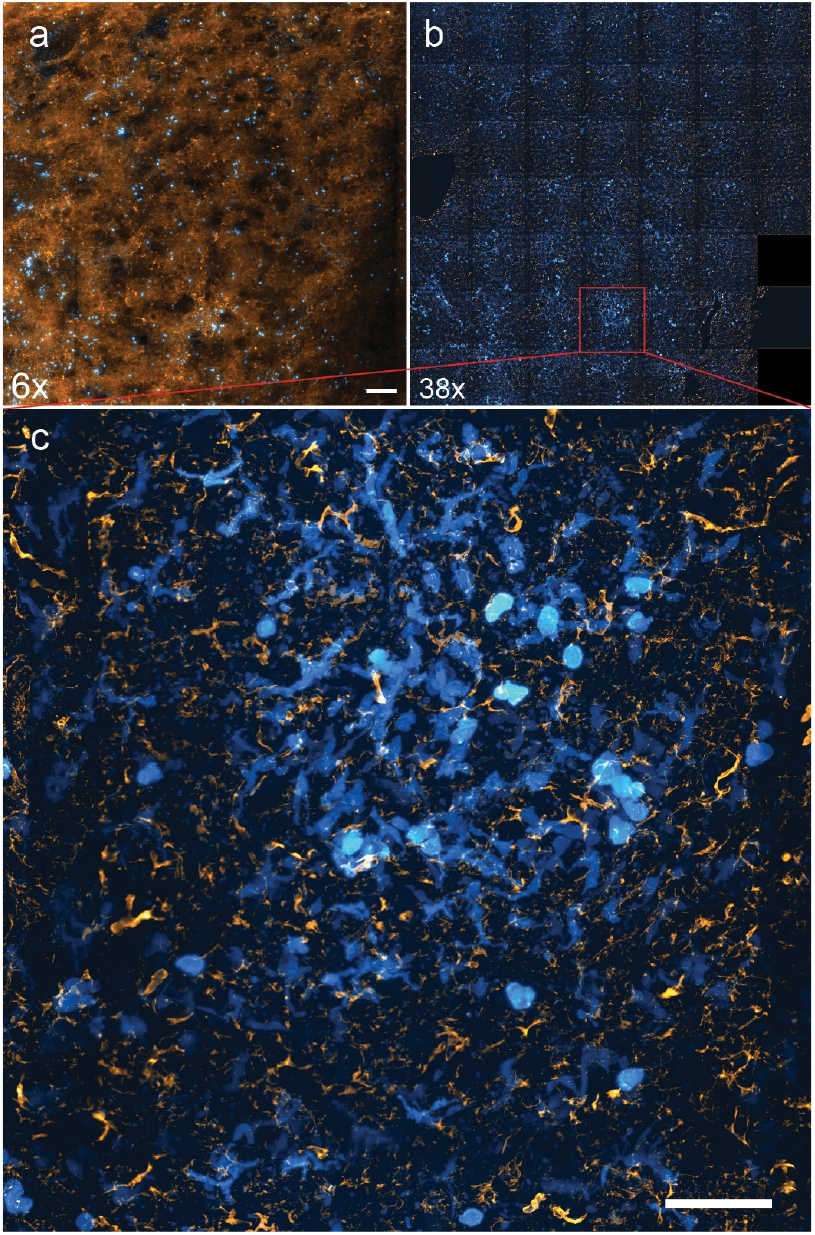
Feature Driven Autonomous Imaging. **a)** Mouse lung sample stained for endomucin (yellow) and platelet factor 4 (PF4), which is alco known as CXCL4 (blue), imaged at 6x magnification with the macroscale module. The tissue boundary was automatically detected and a tiling pattern for high-resolution imaging with the nanoscale module established. b) High-resolution tiled image acquisition. c) A zoomed-in section at 38x magnification is shown at the bottom. Scale bars: A, 6x = 170 μm, 38x = 50 μm.

Understanding the spatial relationship between glomeruli and surrounding nerves is critical for unraveling the mechanisms of innervation’s role in kidney development and physiological regulation. To characterize glomeruli associated with nerves, we developed a workflow where the entire kidney (P0) was volumetrically imaged at 6x magnification with our macroscale module (**Figure 5a**). Candidate glomeruli were automatically detected on-the-fly with a custom segmentation approach (see **Methods**) and interrogated at high resolution with our nanoscale module (**Figures 5b and c**), following the most efficient imaging path. The size of the Z-stack for the nanoscale module was dynamically set based on the maximum feature size identified with the macroscale module. Leveraging this fully automated pipeline, we successfully imaged approximately 839 glomeruli at high-resolution, encompassing over 1 million image slices.

**Figure 5.**
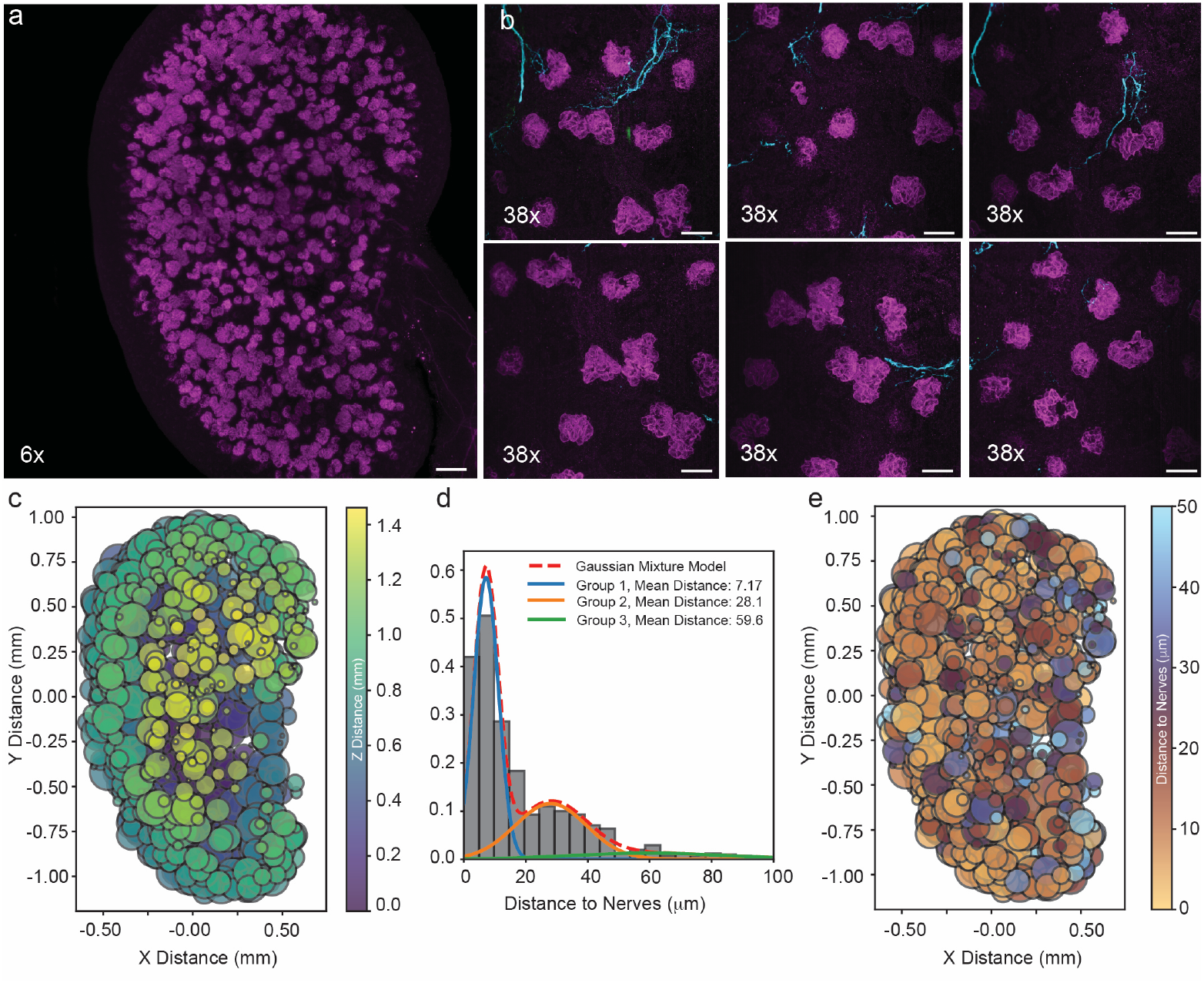
Feature-driven autonomous imaging of glomeruli and innervation in a mouse kidney. **a)** Whole mouse kidney imaged at 6x magnification with the macroscale module, stained for Podocin (magenta), which labels glomeruli, and TUBB3 (cyan), which labels nerves. On-the-fly segmentation was performed immediately after acquisition to identify glomeruli, enabling targeted high-resolution imaging with the nanoscale module. **b)** Representative high-resolution images of glomeruli captured with the nanoscale module, highlighting the diversity of innervation across different regions. **c)** Scatter plot showing the position of glomeruli in X, Y, and Z (color-coded as per the lookup table) and their volumes (represented by the diameter of data points). **d)** Histogram of distances between glomeruli and adjacent nerves. The data was fit to a mixture model of three Gaussian populations with mean distances of 7.17, 28.1, and 59.6 μm. Individual Gaussian populations are shown as solid lines, and the cumulative mixture model as a dashed line. **e)** Scatter plot showing glomeruli position in X and Y, with color-coded distances to adjacent nerves (lookup table) and glomeruli volume (represented by data point diameter).

To further analyze this dataset, we evaluated glomerular volume, position, and proximity to nerves. Of the 839 positions imaged, 835 included glomeruli, demonstrating the robustness of the automated segmentation and targeting workflow. Glomeruli in the high-resolution datasets were detected using a difference of Gaussian filter and an automatic multi-level segmentation. Only the central glomeruli in the volume were evaluated. Scatter plots depicting the position of glomeruli in X, Y, and Z, as well as glomerular volume, revealed the distribution of glomeruli size throughout the entire kidney (**Figure 5c**). As expected, superficial glomeruli were generally smaller due to their early developmental stage compared to more mature glomeruli located deeper in the cortex. Next, we evaluated nerve proximity to glomeruli. Nerves were identified using a Meijering neurite filter. By plotting the distances between glomeruli and nearby nerves as a histogram, three distinct populations of glomeruli emerged, with mean distances to nerves of 7.17, 28.1, and 59.6 microns (**Figure 5d**). The majority of glomeruli had nerve fibers within 50 microns, a distance suggested to be sufficient for signal diffusion to target cells^21^. Those within Group 1 and a mean distance of 7.17 microns likely represent close associations of nerves with Bowman’s capsule and glomerular arterioles^3^. To assess whether there was any spatial organization to these distances, we created a scatter plot where each glomerulus was color-coded based on its distance to the nearest nerve (**Figure 5e**). The superficial glomeruli appeared to have closer associations with nerves which may correlate with the later embryonic onset and expansion of kidney innervation^3^. These analyses highlight the utility of MCT-ASLM for comprehensive imaging workflows, enabling the capture of high-quality data that supports global analyses of tissue architecture. Such capabilities provide critical insights into spatial relationships, such as glomerular innervation patterns, that are essential for understanding kidney development.

## Discussion

A key challenge in biology is understanding how cellular function varies throughout a tissue, influencing local and global architectures and shaping developmental, physiological, and pathological outcomes^2^. MCT-ASLM addresses this by enabling multiscale imaging of large, chemically cleared specimens, linking subcellular structures to tissue-wide organization. Its two complementary imaging arms, the macroscale^15^ and nanoscale modules^22^, are arranged in a 4-axis geometry that facilitates smooth transitions between large field-of-view imaging and high-resolution interrogation. When combined with sample rotation, this design supports volumetric imaging of specimens as large as 20 mm^3^, resolving features as small as ~300 nm. This unique combination of capabilities makes MCT-ASLM an ideal tool for biological studies that demand both comprehensive tissue context and detailed cellular resolution.

Paired with navigate^13^, our intelligent microscope software, MCT-ASLM operates in an intuitive, multi-resolution format that includes human-guided and autonomous modes. Event detection is achieved through adaptable classical computer vision routines, which can be readily modified for diverse specimens without the need for computationally intensive retraining of neural networks. These routines allowed us to implement self-driving features, including automated multi-resolution and multi-module tiling and the detection and interrogation of over 800 glomeruli in a developing kidney. By automating feature identification and imaging, the system eliminates the need for manual intervention, substantially increasing throughput while ensuring high-contrast, high-resolution operation of the microscope. Such smart routines reflect the broader evolution of smart microscopy, which has been used to optimize imaging parameters dynamically^23^, reduce light exposure^24^, adjust imaging resolution^25^, correct optical aberrations^26^, and autonomously detect and interrogate transient or otherwise rare biological events, such as mitochondrial fission events^27^ or Plasmodium-infected cells^28^, respectively. Indeed, our approach aligns with recent advances in smart lattice light-sheet microscopy, which autonomously switched imaging modes in response to detected events, enabling population-level statistics to be captured much faster than through manual approaches^12^.

The benefits of this system extend beyond convenience. Compared with classical imaging approaches, where the entire specimen is imaged, MCT-ASLM’s throughput improves as the feature of interest becomes less frequent. Thus, MCT-ASLM is particularly useful for imaging rare cell populations, such as stem or progenitor cells in tissues like bone marrow, liver, or intestine, which are often identified using combinations of cell surface markers or reporter labeling^29^. Here, we evaluated low-frequency metastatic colonization of bone and lung tissues, which conventionally requires exhaustive specimen-wide imaging or labor-intensive manual histology-based identification. Similarly, we identified sparsely labeled neurons in rat brains. Moving forward, feature identification could be increasingly sophisticated and multi-parametric, leveraging logic gates to evaluate heterotypic cell-cell interactions^30^ or identifying select subsets of cells according to morphological^31^, signaling^32^, or gene expression^33,34^ profiles.

In summary, MCT-ASLM bridges spatial scales, offering a transformative solution for imaging cm-scale tissues while resolving subcellular structures. By integrating smart microscopy principles—such as automated imaging paths, dynamic parameter adjustments, and multi-resolution workflows—it facilitates rapid and precise analysis of large, complex specimens. This capability to identify and interrogate features with high resolution in an efficient, automated manner makes MCT-ASLM an invaluable tool for advancing research across developmental biology, cancer studies, and neuroscience, paving the way for accelerated scientific discoveries.

## Acknowledgements

The authors would like to acknowledge Drs. Reto Fiolka and Gaudenz Danuser for their insightful conversations, and Dr. Dana Reed and Ms. Rebekah Craig for their invaluable administrative support. The authors received generous funding from the National Institutes of Health National Cancer Institute (U54CA268072, P30CA142543, R01CA295997, R01CA238519, T32CA291654), National Institute of General Medical Sciences (RM1GM145399, R35GM142654), National Institute of Neurological Disorders and Stroke (R01NS127900, R21NS128203), National Institute of Diabetes and Digestive Kidney Diseases (R01DK133283, R01DK121014). Additional support was provided by the Trauma Research and Combat Casualty Care Collaborative (175145), Charles Pak Family Cancer & Bone Initiative, Lyda Hill Philanthropies, American Heart Association (24PRE1241584), and Cancer Prevention and Research Institute of Texas (CPRIT) (RR1900371, RP230261, RP240183, RR180014). Z.Z and S.J.M. are CPRIT Scholars. S.J.M. is a Howard Hughes Medical Institute (HHMI) investigator, the Mary McDermott Cook chair in Pediatric Genetics, the Kathryn and Gene Bishop distinguished chair in Pediatric Research and the director of the Hamon Laboratory for Stem Cells and Cancer.

## Author Contributions

J.L., Z.M., and K.M.D. designed and built the microscope. Z.M., K.M.D., and X.W. programmed the software. H.M.B., P-E.Y.N., X.L., B.A.P., S.D.C., Y.X., M.T.I., A.B.A., H.Z., S.J.M., S.L., Z.Z., L.L.O., and K.M.D. developed the labeling and clearing protocols and prepared the samples. K.M.D. and J.L. performed data analysis and prepared the figures. K.M.D. oversaw all aspects of the research and wrote the manuscript. All authors read and approved the manuscript.

## Online Methods

### Microscope Design

The multiscale microscope consists of macroscale and nanoscale imaging modules built in a 4-axis illumination and detection geometry on a Performance Series CleanTop optical table equipped with UltraDamp air vibration isolators. A detailed parts list is provided in **Supplementary Table 1**. Fiber-coupled illumination (NA = 0.11) is provided to each imaging module with a LightHub Ultra (Omicron-Laserage Laserprodukte GmbH) equipped with compact diode pumped semiconductor lasers emitting at 488 (LuxX 488-150, 150 mW), 561 (OBIS LS, 150 mW), and 642 (LuxX 642, 140 mW) nm. Each laser was operated in a mixed analog and digital modulation mode, enabling rapid and independent control of laser power. Switching between the fiber outputs was achieved with a TTL-triggered mirror galvanometer located within the LightHub Ultra.

For imaging, the sample is positioned in the shared chamber for the macroscale and nanoscale modules from above using precision XYZθ (X, L-509-.20DG10; Y, L-509-20DG10; Z, L-509.40DG10; θ, M-060.DG. Physik Instrumente) stages mounted to a gantry. Sample positioning is coordinated with a brushless DC motor controller (C-884.6DC, Physik Instrumente). All analog and digital waveforms are delivered to the microscope via a National Instruments-based PXIe chassis (PXIe-1073, NI) equipped with a PXI-6259 (32 Analog Inputs, 48 Digital I/O, 4 Analog Outputs) and a PXI-6733 (8 Analog Outputs) connected to noise rejecting breakout boxes (BNC-2110) via shielded cables (SHC68-68-EPM). Operation of the microscope was performed with navigate, which is publicly available on GitHub (https://github.com/TheDeanLab/navigate).

All data are saved in accordance with the Open Microscopy Environment (OME) Data Model. For tiff files, all metadata was provided as an OME-XML. For hierarchical file formats, which provide highly parallelized read/write capabilities that are amenable to cloud-based computing, we adopted the OME-Next-Generation File Format (OME-NGFF) and Zarr/N5/HDF5 file formats. The acquisition computer was a Colfax ProEdge SXT9800 Workstation with dual Intel Xeon Silver 4112 CPUs clocked at 2.60 GHz and outfitted with 96 GB of RAM, running on a 64-bit Windows 10 Pro (Version 22H2) operating system. The operating system was installed on an Intel DC P3100 1 TB M.2 NVMe SSD (x4, 22×80), and data saving operations were performed with a Micron 5200 ECO 7.68 TB SSD. Network connectivity was provided by an Intel X550-T2 RJ45 10Gbase-T PCI-E x8 Dual Port 10GbE adapter.

### Macroscale Module Design

The macroscale imaging module is a single-sided illumination variant of mesoSPIM^15^. Briefly, a fiber from the laser launch is connected to a Thorlabs-based cage system with a fiber adapter (SM1FCA, Thorlabs). Light emanating from the fiber is collimated with a 75 mm achromatic doublet (AC254-075-A-ML, Thorlabs), resulting in a beam diameter of 16.5 mm. To introduce defocus into the beam, the light is passed through an electrotunable lens (EL-16-40-TC-VIS-5D-1-C, Optotune), which is relayed with two achromatic doublets (AC254-080-A-ML, Thorlabs) to a 10 mm galvanometer (GVS211, Thorlabs) that is located at the back pupil of modified DSLR lens (AF-S Nikkor 50 mm f/1.4G, Nikon). Illumination is performed in a digitally scanned^35^ Axially Swept Light-Sheet Microscopy (ASLM)^14^ format. Specifically, a 2D Gaussian beam is laterally scanned to create a virtual (or ‘digital’) sheet of light while an electrotunable lens scans the beam in its propagation direction synchronously with the rolling shutter readout of the camera^15^. Translation of the entire illumination path is achieved with a large-area stage (TBB0606, Thorlabs).

Fluorescence detection is achieved with a research grade macroscope (MVX10, Olympus America, Inc.), a 1X imaging objective (MVXPLAPO1X, NA 0.25, Olympus America, Inc.), a tube lens (MVX-TLU-VX10, Olympus America, Inc.), and a scientific Complementary Metal Oxide Semiconductor (sCMOS) camera (ORCA-Flash 4.0 v3, Hamamatsu Photonics). To eliminate unwanted light, a 10-position filter wheel (LB10-W33 and LB10-3, Sutter Instruments) equipped with green (FF01-515/30-32, Semrock), red (FF01-595/31-32, Semrock), and far-red emission filters (BLP01-647R-25, Semrock) is placed in the infinity space between the macroscope and its tube lens. A motorized servo (Dynamixel MX-28R, Robotis) adjusts the magnification of the macroscope between 0.63x and 6.3x, thereby providing an FOV of 21.1 to 2.1 mm in X and Y. Care was taken to ensure that the image did not translate laterally during changes in magnification, requiring precise positioning of the camera and the tube lens relative to the MVX10 macroscope, as well as tip/tilt alignment of the sample chamber. Both the illumination and detection paths were introduced into the shared sample chamber through 50.8 mm diameter, uncoated, 1 mm thick optical windows (Edmund #11-751), which were sealed to the aluminum chamber using low-strain UV-initiated adhesive (Norland NOA65). The entire macroscale detection path is mounted on a long working distance precision stage (M-406.4PD, Physik Instrumente), controlled with the C-884.6DC, Physik Instrumente.

### Nanoscale Module Design

The nanoscale imaging module consists of an improved variant of Cleared Tissue ASLM^36^ (CT-ASLM) that is built in a compact and cage system-based architecture with a beam height of ~94 mm. Here, the second fiber from the laser launch is connected to a fiber adapter (SM1FCA, Thorlabs) and collimated with an achromatic doublet (AC254-50-A-ML, Thorlabs). Polarization of the beam (diameter = 11 mm) is adjusted with a half-wave plate (10RP52-1B, Newport) positioned in a rotational mount (CRM1T, Thorlabs). Thereafter, the light is focused with an achromatic cylindrical lens (ACY-254-50-A, Thorlabs) onto a 4 kHz resonant galvanometer (CRS4KHz, Novanta), recollimated with an achromatic doublet (AC254-100-A-ML, Thorlabs), passed through a polarizing beam splitter (CCM1-PBS251/M, Thorlabs) and a quarter wave plate (AQWP3, Bolder Vision) before ultimately being delivered to an air immersion remote focusing objective (CFI60 Super Plan Fluor LWD 20X, NA 0.7, Nikon Instruments). To back-reflect the light, a mirror is placed at the nominal focal plane of the remote focusing objective and scanning of the mirror is performed with a one-dimensional translatory voice coil (LFA2004, Equipment Solutions), controlled with a servo (SCA814, Equipment Solutions). The back-reflected light subsequently makes a second pass through the quarter wave plate, is reflected off the polarizing beam splitter, and relayed to the back pupil of the multi-immersion illumination objective (54-12-8, Applied Scientific Instrumentation) with two achromatic doublets (AC254-125-A-ML and AC254-150-A-ML, Thorlabs). Operation of the resonant galvanometer, which is conjugate to the specimen, reduces shadow artifacts by rapidly pivoting the illumination^37^. For detection, the emitted fluorescence is captured with an identical multi-immersion objective (54-12-8, Applied Scientific Instrumentation), and focused with an achromatic doublet (AC508-300-A, Thorlabs) onto back-thinned sCMOS camera (ORCA-Fusion BT, Hamamatsu Photonics). Unwanted light is rejected with a 10-position filter wheel (LB10-W33 and LB10-3, Sutter Instruments) equipped with green (FF01-515/30-32, Senrock), red (FF01-595/31-32, Semrock), and far-red emission filters (BLP01-647R/31-32, Semrock), which is located between the achromatic doublet and the camera. Compared to previously published variants of CT-ASLM, an upgraded aberration-free remote focusing unit increases the FOV from ~327 to ~417 microns in the X and Y dimensions. To isolate the system from vibrations arising from the voice coil, the entire imaging system was built on an 18” x 24” x 2.4” breadboard (B1824F, Thorlabs) with 95 mm optical rails for facile one-dimensional positioning of the microscope.

### Analysis of Microscope Resolution

To evaluate the resolution of the imaging system, we prepared bead samples at two different concentrations. For the high-resolution setup, 200 nm yellow-green beads (17151-10, Polysciences) were diluted 1:100 in a 2% agarose solution, while a 1:500 dilution was used for the low-resolution system. The agarose solution was liquefied by heating in a microwave, then thoroughly mixed with the bead solution. The hot mixture was poured into a custom mold with a 3D-printed sample holder, forming a five-sided cube. Once the agarose solidified, the sample holder was carefully detached, leaving the bead-filled agarose cube securely attached. Bead data were analyzed using a MATLAB pipeline to calculate Full-Width Half-Maximum (FWHM) values. Beads were first identified via thresholding, and their centroids were located using regionprops3. Beads near image edges and those with a coefficient of determination (R^2^) below 0.9 were excluded. For the remaining beads, line profiles along the X, Y, and Z axes were extracted and fitted with Gaussian functions. Sigma values from these fits were converted into FWHM measurements, which were then exported as CSV files for further analysis in Python.

### Vertebrate Animals

All procedures involving vertebrate specimens were conducted in accordance with the National Institutes of Health Guide for the Care and Use of Laboratory Animals and received approval from the Institutional Animal Care and Use Committees at the University of North Carolina at Chapel Hill, the University of Texas Southwestern Medical Center, and the China Institute for Brain Research.

### Stereotaxic AAV Injection

Stereotaxic AAV injection was performed on SD rat of 12 weeks age. Two types of viruses, AAV2/9-hsyn-Cre (titer 13 × 1012, diluted by 10000 folds) and AAV2/9-EF1α-DIO-mScarlet (titer 8.7 × 1012), were mixed at 1:1 ratio and injected into the M2 region (anteroposterior, 5.16 mm from bregma; mediolateral, 2.5 mm; dorsoventral, −3.0 mm) or the CG1 region (anteroposterior, −0.4 mm from bregma; mediolateral, 1.56 mm; dorsoventral, −3.0 mm). After injection, the micropipette was left in place for 10 minutes before withdrawn. Rats were allowed to recover for 6 weeks before sacrificing.

### Benzyl Alcohol Benzyl Benzoate (BABB) Clearing Protocol

Lungs and livers were cleared using a modified BABB protocol. All steps were performed with gentle rotation, and samples were protected from light. Freshly collected tissues were fixed in 4% PFA at 4°C for less than 24 hours. After fixation, tissues were washed in PBS with 0.02% sodium azide three times for 2 hours each, followed by overnight incubation at 4°C. Tissues were then cut into ~1-2 mm slices using a matrix and placed in individual 1.5 mL tubes. Pre-treatment involved incubating the samples in 25% QUADROL at 37°C with shaking, and non-perfused tissues were refreshed until the supernatant was clear. Perfused samples were incubated overnight in 25% Quadrol. Following Quadrol treatment, tissues were washed with PBS and incubated in blocking buffer (PBS + 0.5% NP40, 10% DMSO, 5% serum, 0.5% Triton X-100) for 1 day at room temperature. Samples were then incubated in primary antibody diluted in staining buffer for 72 hours at room temperature, shielded from light. After primary labeling, tissues were washed in Wash Buffer (PBS + 0.5% NP40, 10% DMSO) with buffer changes every 2 hours for 6 hours. The samples were post-fixed overnight in 4% PFA. Following post-fixation, tissues were washed with PBS and incubated in secondary antibody in staining buffer for 72 hours at room temperature, protected from light. After secondary labeling, tissues were washed with Wash Buffer every 2 hours for 6 hours and left in Wash Buffer overnight. The next day, Wash Buffer was refreshed. For nuclear staining, tissues were incubated in a dye such as SYTOX Green 488 (1 µM) for 48 hours at room temperature. After staining, tissues were washed twice with PBS for 15 minutes and dehydrated in a methanol gradient (25%, 50%, 75%) for at least 1 hour per step. The samples were then placed in 100% methanol for 45 minutes, refreshed, and rotated for an additional 45 minutes. After methanol dehydration, the tissues were delipidated with two 30–45-minute washes in dichloromethane (DCM). Samples were then incubated in fresh BABB (1:2 benzyl alcohol: benzyl benzoate) for tissue clearing, with three buffer changes. The cleared samples were incubated overnight in fresh BABB at room temperature, protected from light. All BABB solutions were prepared by placing 45mL of BABB in a 50mL conical with 5g of activated aluminum oxide (to remove peroxides), gently agitating the solution for at least 1 hour at room temperature and centrifuging the tube at 3000rpm for 10 mins to pellet the aluminum oxide.

### Modified iDISCO Clearing Protocol

For tissue clearing of embryonic and postnatal mouse kidneys or urogenital systems (P07), we adapted a modified version of the iDISCO protocol^38^ for whole-mount immunolabeling as described previously^3^. All steps were performed on a rotating shaker with the specimen protected from light. Pre-fixed tissues were first incubated in 1 mL of blocking solution (ice-cold PBS with 10% v/v Heat Inactivated Sheep Serum (ThermoFisher, 16070096) and 0.5% v/v Triton X-100) for 1 hour at room temperature. Blocked samples were incubated in 1 ml of primary antibody diluted in the blocking solution for 7 days at 4°C. Antibodies and concentrations used are provided in **Table S5**. After primary labeling, tissues were washed for 24 hours at room temperature in 1 ml of wash buffer (PBS with 0.25% Triton X-100), with the wash solution replaced every hour. Thereafter, the samples were incubated in 1 ml of secondary antibody diluted 1:1000 in the blocking solution for 3.5 days at 4°C. Post-secondary labeled tissues were washed for 24 hours using the same wash protocol and embedded in a ~5 mm x 5 mm agarose block (Fisher Science, BP160-500). For tissue clearing, agarose blocks were sequentially dehydrated in a methanol-water series (25%, 50%, 75%, and 100%) at room temperature, 1 hour per step. Dehydrated blocks were incubated in a 66% dichloromethane (DCM) (Sigma, 270997)/33% methanol mixture for 3 hours at room temperature, followed by two 15-minute washes in 100% DCM. Finally, the samples were cleared by incubation in 100% dibenzyl ether (Sigma, 33630) at room temperature until transparent (5-7 hours). Samples were stored in dibenzyl ether until imaging.

### Tissue Clearing with the TESOS Method

The TESOS method (transparent embedding solvent system) was performed with slight modifications based on previous publication^39,40^. Adult rats were perfused, and brains were fixed in 4% PFA at 4 °C for 24 hours. Samples were decolorized with a 25% (w/v in H2O) Quadrol (Sigma-Aldrich, 122262) + 10% tert-Butanol solution (tB, Sigma-Aldrich, 360538) at 37 °C for four days. Next, rat brain samples were immersed in gradient delipidation solutions at 37 °C for two day per concentration: 30% tB solution (v/v in H2O), 50% tB solution (v/v in H2O), and 70% tB solution (v/v in H2O). Each gradient delipidation solution was supplemented with a final concentration of 3% (w/v) Quadrol to maintain the pH over 9.0. Following this, rat brains were dehydrated in a tB-Q dehydration medium consisting of 70% (v/v) tB, and 30% (w/v) Quadrol at 37 °C for four days. Finally, the brains were immersed in TESOS clearing medium (refractive index R.I. 1.55) at 37 °C until full transparency, which typically took around two to three days. The TESOS clearing medium comprised of 53% (v/v) benzyl benzoate (BB) (Sigma Aldrich W2131802) and 32% (v/v) bisphenol-A ethoxylate diacrylate Mn 468 (BED468) (Sigma-Aldrich 413550) “10% EGDMA (v/v) (Sigma Aldrich, 335681) with 5% (w/v) Quadrol. Cleared samples were preserved in BB-BED solution at 4°C. BB-BED solution was also used as the imaging medium.

### Segmentation of Glomeruli

To enable rapid and automated segmentation of 3D volumetric data, we implemented a pipeline leveraging Dask (https://www.dask.org/) for parallelized processing and efficient handling of large datasets. This segmentation process was performed automatically on the microscope immediately after imaging with the macroscale module, ensuring seamless integration into the imaging workflow. While most of navigate’s features perform image analysis on data stored in RAM, the 3D object detection feature was specifically designed to load data directly from the hard drive, making it ideal for large datasets; we recommend saving data in N5 or Zarr file formats, which support highly parallel access for optimal performance. If the raw image is saved as a TIFF file, it is first loaded and converted into a Dask array to accelerate processing by distributing computation across available CPU or GPU resources. For N5 or Zarr file formats, this conversion step is not necessary, as the data can be directly accessed in a parallelized manner. To minimize the influence of outlier intensities, voxel values above the 99.99th percentile are capped at 100. This adjustment ensures that exceptionally bright regions do not skew the subsequent filtering and thresholding steps. High-pass filtering is performed using a 3D Gaussian filter with a FWHM of 10 voxels, while low-pass filtering uses a 3D Gaussian filter with a FWHM of 20 voxels. Subtraction of the high- and low-pass filtered images creates a Difference of Gaussian image, enhancing intermediate spatial frequencies associated with objects of interest. The difference image is down sampled, and multi-level Otsu thresholding is applied to identify intensity thresholds for separating the data into distinct classes. The second threshold is used to binarize the full-resolution difference image, isolating regions of interest automatically. A distance transform is applied to the binary image, calculating the distance of each voxel to the nearest background voxel. Voxels with distances greater than or equal to 7 voxels are retained, producing a refined binary mask that isolates well-separated objects. The refined binary mask is processed with connected component labeling to assign unique labels to each distinct object. Background regions are excluded, and the number of unique objects is recorded. The labeled image, along with the locations of all identified objects for subsequent interrogation with the target imaging mode, is saved for future evaluation. By utilizing Dask to parallelize computational tasks, this pipeline provides a versatile framework for segmentation routines that can be readily adjusted to accommodate different object types. Dask methods such as *map_overlap* and *map_blocks* enable efficient processing of data in a chunk-wise format while mitigating edge artifacts.

### Spatial Analysis of Glomeruli-Nerve Proximity

Glomeruli in the high-resolution datasets were detected using a 3D difference of Gaussian filter, followed by automatic multi-level segmentation. For each imaged volume, the labeled glomeruli were analyzed using region properties, where regions with fewer than 100,000 voxels were excluded to eliminate small, non-glomerular structures. The centroid of each glomerulus was determined, and the glomerulus closest to the center of the imaging volume was identified for further analysis. Nerves were segmented using the Meijering filter, applied slice-by-slice in the XY plane across the 3D dataset. The filtered nerve volumes were thresholded to generate binary masks, which were subsequently used for computing the Euclidean distance transform. Both glomeruli and nerve detection workflows were executed on a distributed computing cluster using ‘dask_jobqueue.slurm.SLURMRunner’, employing 6 nodes with 72 threads and 512 GB of RAM per node. This parallelized approach allowed for efficient processing of the large 3D datasets, enabling rapid and scalable segmentation and analysis. To quantify the spatial relationship between glomeruli and nerves, the distance from the centroid of each glomerulus to the nearest nerve structure was calculated using a distance transform.

### Data Stitching and Fusion

For experiments involving automated tiling, all imaging data were saved as an HDF5, N5, or OME-Zarr file format, along with a BigDataViewer-compatible XML file defining the dataset. These file formats enable efficient handling of large, high-resolution datasets and are optimized for hierarchical storage and parallel access. The inclusion of stage coordinates and camera parameters (e.g., pixel size, magnification, etc.) during dataset generation places tiles into their proper spatial context and significantly accelerates the stitching process and reduces the RAM overhead by limiting comparisons to overlapping regions. Both the Fiji plugin and the standalone BigStitcher-Spark implementation were utilized, the latter of which is particularly valuable for large datasets.

### Deconvolution

Deconvolution of the acquired datasets was performed using the PyPetaKit5D library, a Python-based wrapper for PetaKit5D^16^. Optical transfer function Masked Wiener (OMW) deconvolution was performed with two iterations, Hann window bounds of 0.8 and 1 to control the optical transfer function apodization, a Wiener parameter of 0.005, an optical transfer function cumulative threshold of 0.6, a background intensity of 100, with no edge erosion applied during processing.

