## Supplemental Materials for "Feature-Driven Whole-Tissue Imaging with Subcellular Resolution"

### Supplementary Notes

#### Supplementary Note 1 - Switching Between Macroscale and Nanoscale Imaging Modules

Navigate is a Python-based software package designed to enable seamless configuration and operation of multiple imaging systems. It supports the use of shared or dedicated hardware devices for controlling individual microscopes. For MCT-ASLM, a deliberate displacement between the macroscale and nanoscale imaging positions was introduced to address two key considerations. First, imaging deep into the large chamber used in MCT-ASLM introduces minor but non-negligible amounts of spherical aberration in the macroscale detection arm, which can reduce resolution and contrast. Second, adopting a more spread-out geometry simplifies the introduction and manipulation of samples within the imaging chamber. The displacement between the two imaging systems was precisely determined by aligning the tip of a sharp needle-like point in the center of each microscope's field of view and calculating the offsets required to align the imaging positions. These offsets are specified for the X, Y, Z, F (focus), and θ (rotation) axes and are included in the configuration file for MCT-ASLM. Navigate automatically applies these offsets when switching between imaging modes, whether done manually by the user or through programmed features. Thus, with navigate, switching between the macroscale and nanoscale modules is readily achieved while maintaining optimal sample positioning and imaging performance.

#### Supplemental Note 2 – Design Considerations for MCT-ASLM

The design of MCT-ASLM incorporates several subtle yet important considerations, particularly regarding the mitigation of vibrations, which can significantly impact imaging performance on the nanoscale module. These design choices provide insights that could inform future implementations of multiscale imaging systems. To achieve a compact footprint for the nanoscale module, Thorlabs cage components were utilized. However, the narrow metallic rods in these components were found to readily transmit vibrations in an undamped manner. Additionally, the large gantry supporting the 4D sample positioning stages was insufficiently stiff, amplifying vibrations instead of suppressing them. While these issues did not affect the macroscale module, vibrations became evident during imaging with the nanoscale module. To address these challenges, several strategies were employed. First, all but two vibration sources were removed from the optical table: the actively cooled cameras and the remote focusing voice coil. Upon investigation with an accelerometer, the cameras were found to produce minimal problematic vibrations. And in our experience, switching from air-cooled to water-cooled cameras only replaces a narrow-frequency vibration source with a broader but lower amplitude vibration source. To mitigate vibrations from the voice coil, two measures were implemented. The voice coil was securely mounted on a vibration-damping sandwich mount (94955K31, McMaster-Carr) with a durometer of 50A. This configuration ensured that vibrations originating from the voice coil were significantly attenuated before reaching the optical table where it could affect other components. Additionally, the nanoscale module was physically separated from the voice coil and placed on an elevated breadboard to provide an extra layer of vibration damping. To simplify alignment and positioning, the nanoscale module was mounted on a rail system, allowing precise adjustments without compromising stability. These measures collectively reduced vibration transmission and improved imaging performance on the nanoscale module. Future iterations of MCT-ASLM will likely replace the large gantries and cage components with more rigid and vibration-resistant designs to further enhance stability and performance.

#### Supplementary Note 3 – Navigate Features

*Automatic Adjustment of Imaging Parameters.*

When switching between imaging systems (e.g., the macroscale and nanoscale modules) and magnifications (e.g., between the 1x and 6x macroscale imaging modes), navigate can automatically adjust various microscope parameters as needed. For instance, the *SetCameraParameters* feature allows for changes in sensor mode, readout direction, and the width of the rolling shutter. Similarly, *UpdateExperimentSetting* can be used to modify imaging parameters such as laser settings (wavelength, intensity), emission filter, or number of imaging channels. This feature also ensures that metadata for image acquisition is updated in real time, maintaining accuracy in the recorded data.

*Automated Multiscale Tiling.*

Navigate uses the *VolumeSearch* feature, which enables the automatic detection of tissue boundaries followed by targeted higher-resolution tiling within the detected regions. As previously published, this feature allows seamless transitions between magnifications in a single imaging system, or as demonstrated here, between imaging systems. The protocol begins with specifying the target imaging system (e.g., macroscale or nanoscale) and magnification (e.g., 1x, 6x, or 38x). Navigate assumes that the microscope is initially operating at the starting resolution and magnification. The system first images the specimen at the starting resolution and segments the acquired data—using an Otsu threshold for segmentation in this case. More advanced segmentation algorithms can be integrated into navigate through its graphical user interface. Based on the segmentation, navigate generates a list of positions to be imaged in the target operating mode, incorporating user-defined overlap between adjacent image tiles for seamless reconstruction.

*Automated 3D Object Detection and Imaging.*

To enable fully automatic 3D object detection and imaging, we implemented a featured called VolumeSearch3D that segments and maps structures in 3D datasets, followed by the automatic acquisition of high-resolution data in a multi-position format. The protocol begins by specifying parameters such as the target imaging system (e.g., macroscale or nanoscale), target magnification level, and the desired Z-step size. The system first acquires a Z-stack of the sample with the current imaging system and magnification (e.g., macroscale and 6x, respectively). Segmentation is then performed on the dataset using a user-provided function, which in this example was a 3D Difference of Gaussian particle detection algorithm. Positions for imaging the segmented objects are then calculated based on the offsets between the current and target imaging systems. The positions, along with the corresponding Z-ranges, are saved for subsequent high-resolution imaging. The size of the Z-stack at the target imaging resolution is determined by the largest feature size identified during segmentation, with an additional user-specified padding, typically set to 5 microns; for example, if the largest feature spans 50 microns in Z, a Z-stack of 60 microns is acquired to ensure complete coverage. Once the positions are mapped, the microscope transitions to the target imaging system and magnification, automatically updating parameters such as camera settings, waveform parameters, and imaging channels. High-resolution 3D data is then acquired for each identified object. All positions and labeled object indices are saved in a text file for reproducibility and further analysis.

**Supplemental Figures**

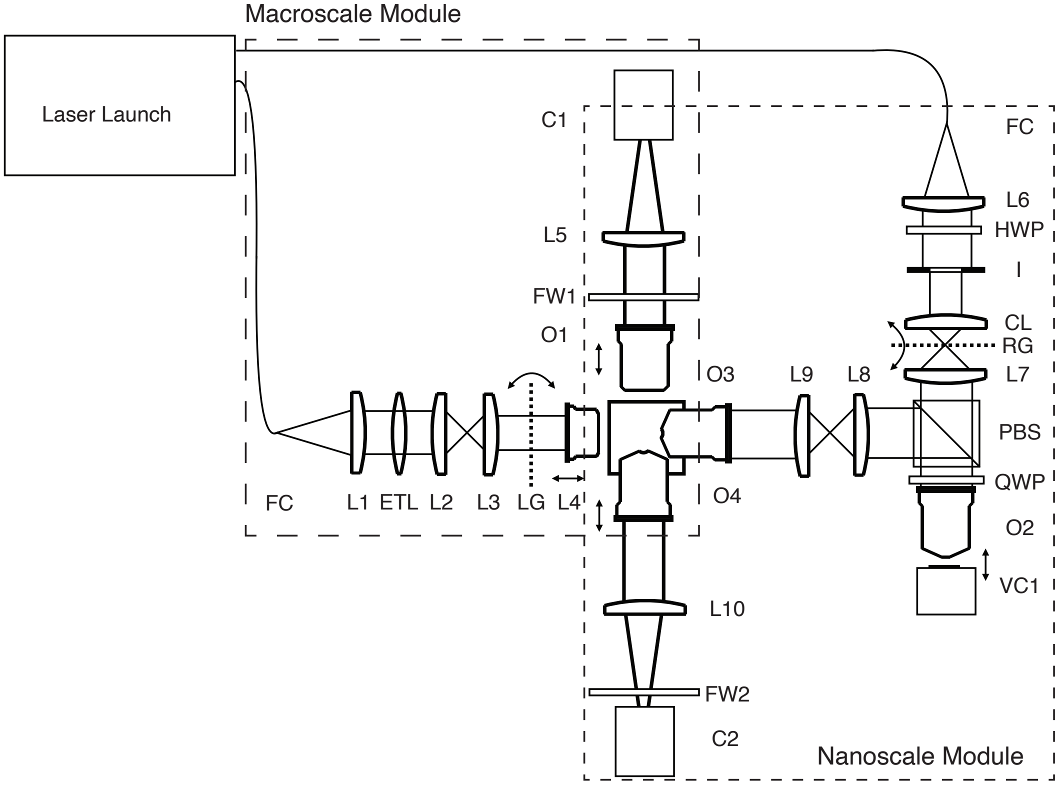

**Figure S1.** MCT-ASLM consists of a macroscale (large dashes) and nanoscale (small dashes) modules that share a common laser launch. The macroscale module includes an FC/APC fiber coupler (FC), 75 mm achromatic doublet (L1), electrotunable lens (ETL), 80 mm achromatic doublets (L2 and L3), linear galvo (LG), a digital single-lens reflex lens (L4), a 1x imaging objective (O1), filter wheel (FW1), tube lens (L5), and a scientific CMOS camera (C1). The nanoscale module includes an FC/APC fiber coupler (FC), 50 mm achromatic doublet (L6), half-wave plate (HWP), iris (I), 50 mm cylindrical lens (CL), resonant galvo (RG), 100 mm achromatic doublet (L7), polarizing beam splitter (PBS), quarter wave plate (QWP), remote focusing objective (O2), voice coil (VC1), 125 mm achromatic doublet (L8), 150 mm achromatic doublet (L2), illumination objective (O3), detection objective (O4), 300 mm achromatic doublet (L10), filter wheel (FW2), scientific CMOS camera (C2).

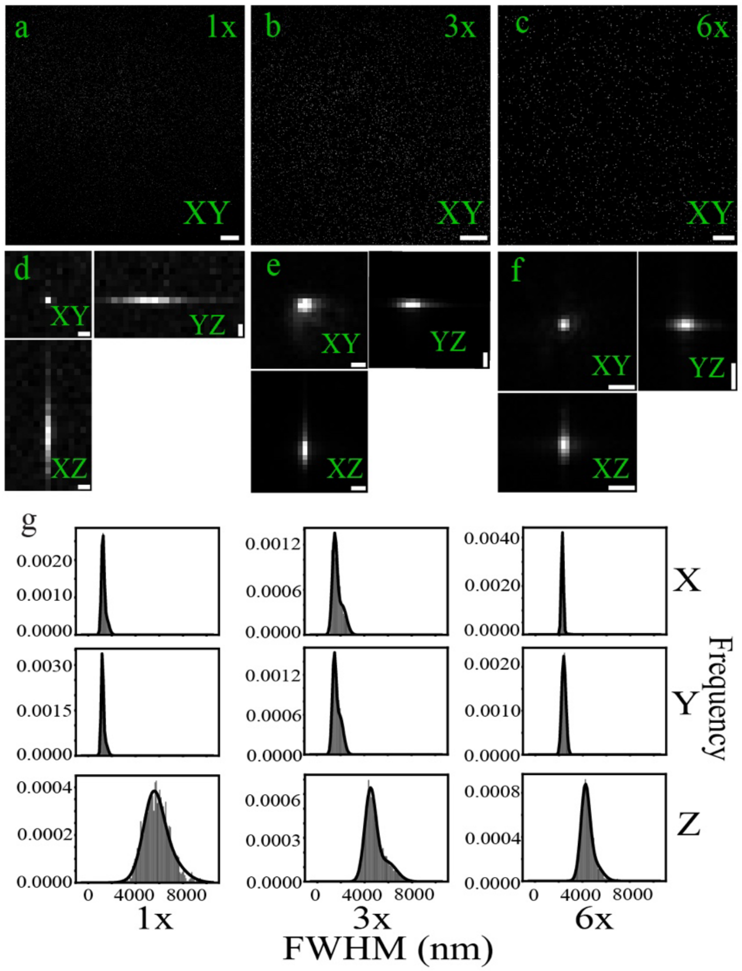

**Figure S2.** Resolution of MCT-ASLM's macroscale module in aqueous contexts. **a-c)** Maximum intensity projections of 1 µm beads in agarose imaged at 1x, 3x, and 6x magnification, respectively. **d-f)** Maximum intensity projections of an isolated 1 µm bead in XY, XZ, and YZ, imaged at 1X, 3X, and 6X magnification, respectively. g**)** Histograms of the Full-Width Half-Maximum (FWHM) of 1 µm beads, imaged at 1x, 3x, and 6x magnification, respectively. Resolution values for each dimension and magnification were binned and fit with a Gaussian mixture model consisting of two populations, and the smaller FWHM population is reported. Mean FWHM values in X, Y, and Z: 1x magnification, 1.2, 1.2, 5.5 microns (N=1599); 3x magnification, 1.5, 1.5, 5.2 microns (N=6651); 6x magnification, 2.3, 2.5, 4.2 microns (N=1807). In the lateral dimensions (e.g., X & Y), only the 6x magnification achieves Nyquist sampling. In the axial dimension (e.g., Z), all reported values are Nyquist sampled. Scale bars: a = 1000 µm; b = 500 µm; c = 200 µm; d = 2 µm; e = 5 µm; f = 10 µm. Results are summarized in Table S3.

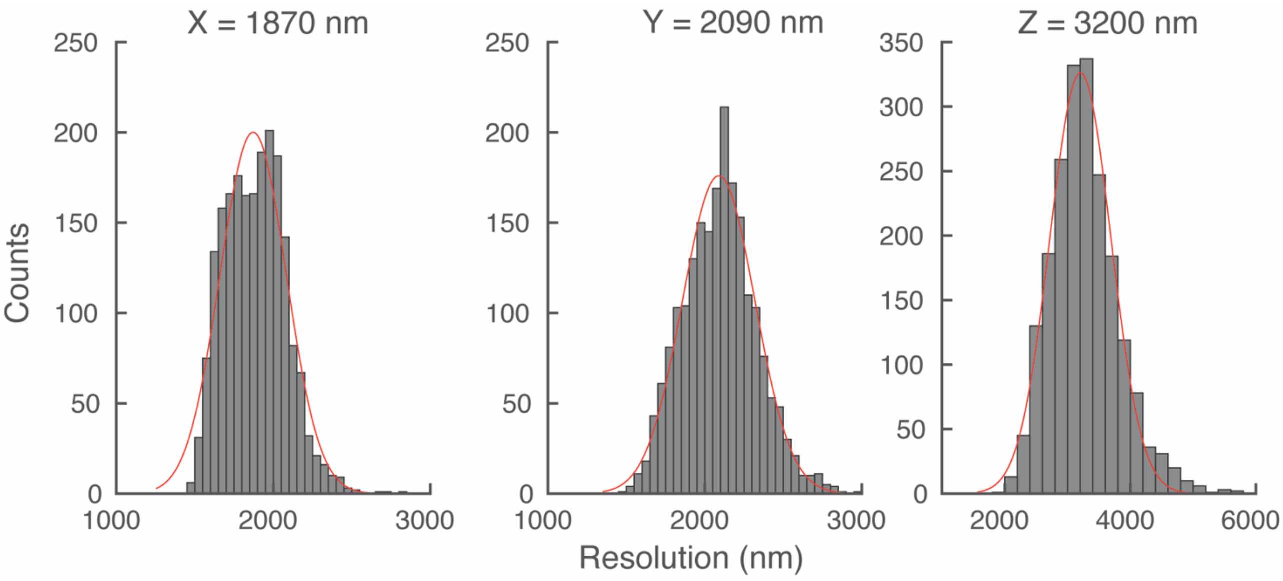

**Figure S3.** Resolution of MCT-ASLM's macroscale module in aqueous contexts after deconvolution. Data were deconvolved with two iterations of the optical transfer function masked Wiener deconvolution method. Beads with a goodness of fit ($R^2$) value greater than 0.9 were included in the analysis. The histogram shows the distribution of full-width half-maximum values for beads imaged at 6x magnification using the macroscale module.

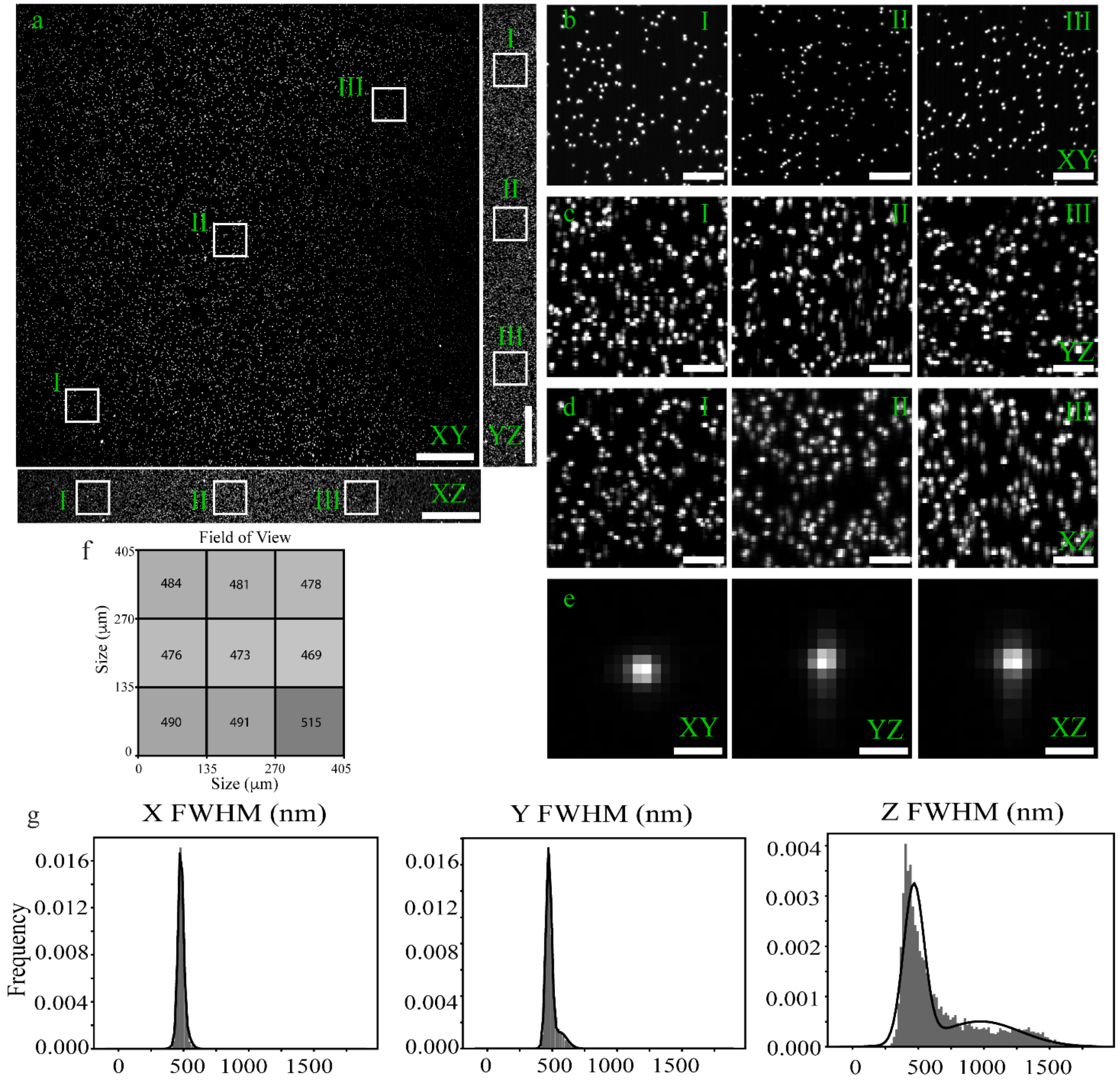

**Figure S4.** Resolution of MCT-ASLM's nanoscale module in aqueous contexts. **a)** Maximum intensity projections of 200 nm beads in agarose. Image volume is 407 µm x 407 µm x 47 µm. Voxel size 198.8 nm x 198.8 nm x 200 nm. **b-d)** Zoomed-in XY, XZ, and YZ maximum intensity projections of 200 nm beads from boxed regions in **a**, respectively. **e)** Maximum intensity projections of an isolated 200 nm bead in XY, YZ, and XZ, respectively. **f)** Heatmap of the lateral FWHM of 200 nm beads across a 407 µm x 407 µm camera chip. **G)** Histograms of the FWHM of 200 nm beads in the X, Y, and Z dimensions. The number of analyzed beads is 8478. Scale bars: a = 50 µm; b, c, d = 10 µm; e = 1 µm.

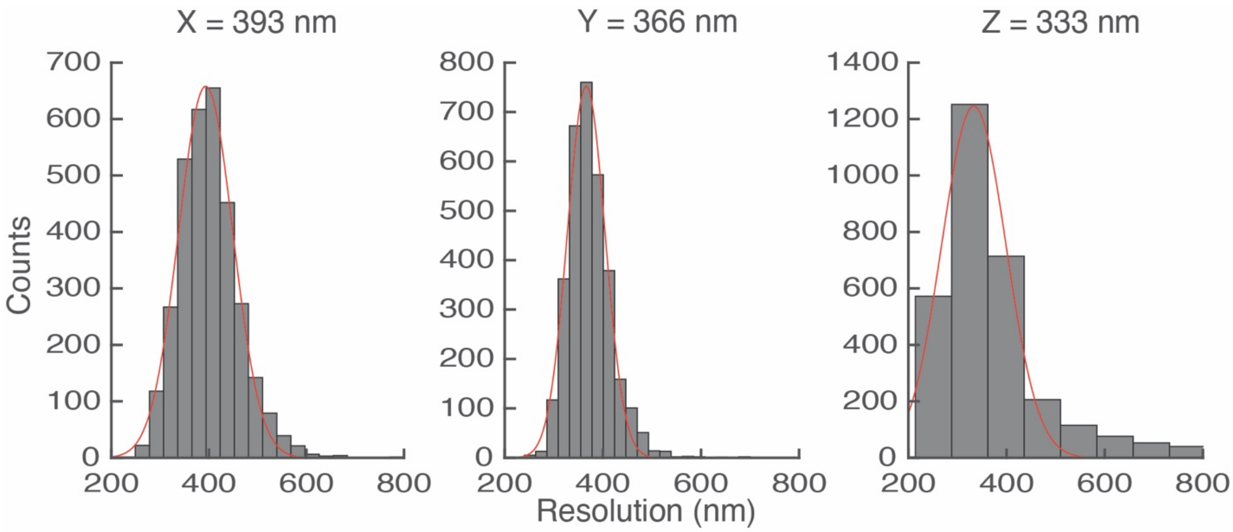

**Figure S5.** Resolution of nanoscale module after deconvolution in aqueous contexts. Data were deconvolved with two iterations of the optical transfer function masked Wiener deconvolution method. Beads with a goodness of fit ($R^2$) value greater than 0.9 were included in the analysis. The histogram shows the distribution of full-width half-maximum values for 200 nm beads.

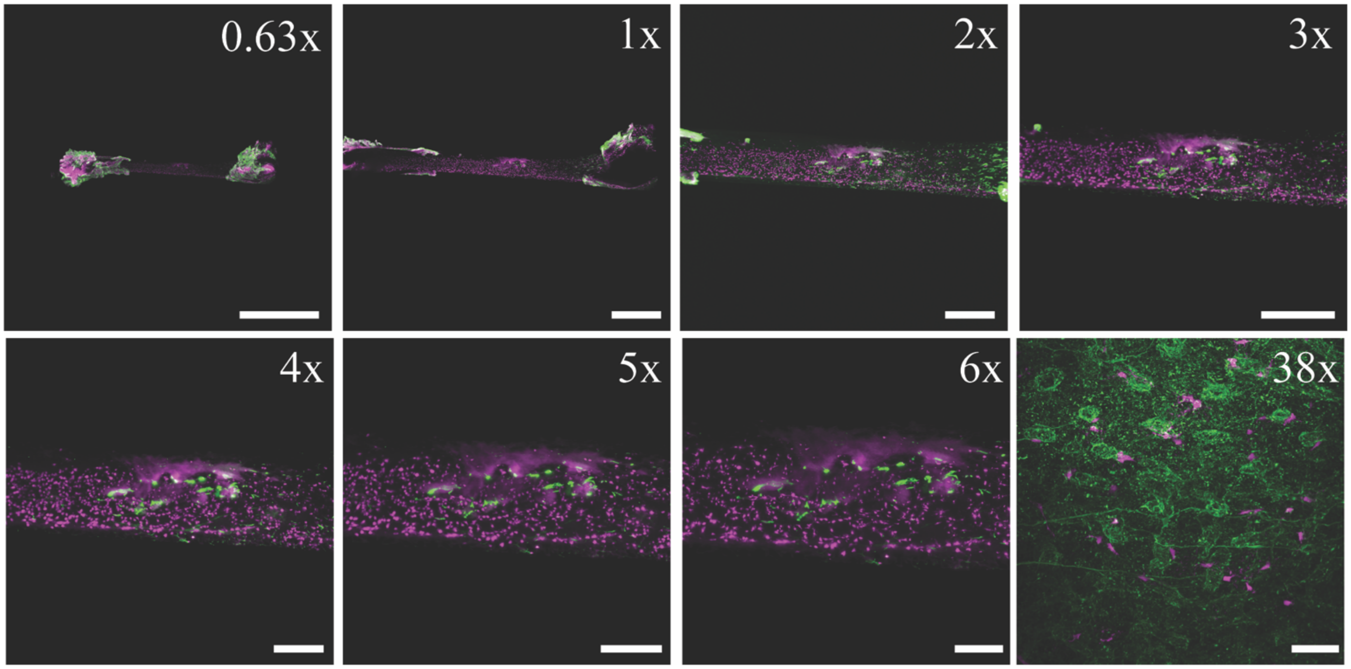

**Figure S6.** BABB cleared mouse bone marrow plug. Sample was imaged with 0.63x, 1x, 2x, 3x, 4x, 5x, 6x, and 38x magnification, respectively. Scale bar: 0.63x = 5000 μm; 1x = 2000 μm; 2x = 1000 μm; 3x = 1000 μm; 4x = 500 μm; 5x = 500 μm; 6x = 300 μm; 38x = 50 μm.

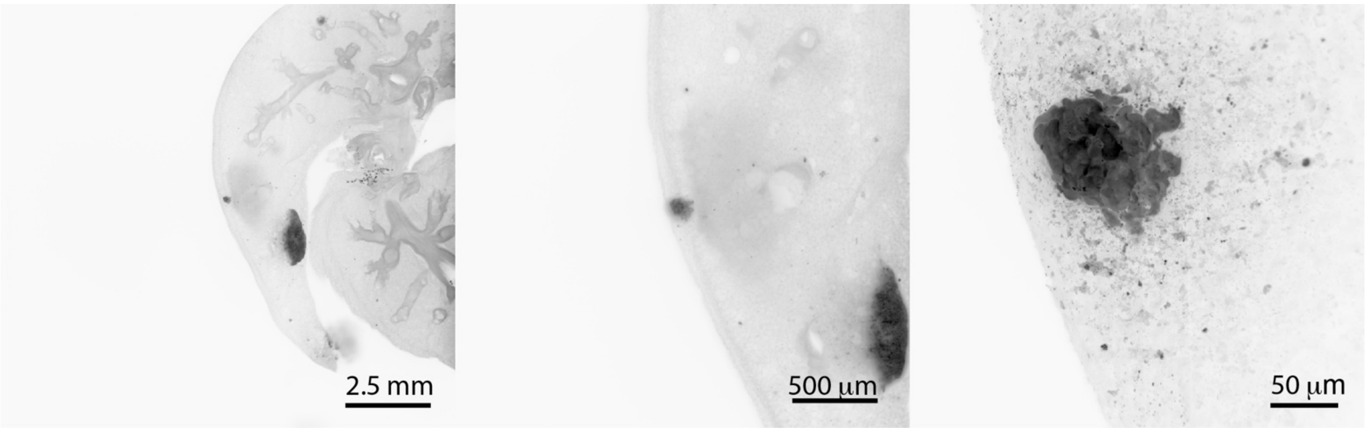

**Figure S7.** BABB cleared mouse lung. Metastatic melanoma was stained with anti-RFP, and the image is inverted to show the tissue architecture. Sample was imaged at 0.63x, 6x, and 38x magnification, from left to right, respectively.

### Supplementary Tables

| Component | Company | Product | Qty |
|---|---|---|---|
| **Common** | | | |
| Laser Launch | Omicron | LightHub Ultra-007B | 1 |
| 488 nm Laser | Omicron | LuxX 488-150 | 1 |
| 561 nm Laser | Coherent | OBIS LS 561-150 | 1 |
| 642 nm Laser | Omicron | LuxX 642-140 | 1 |
| DAQ Chassis | NI | PXIe-1073 | 1 |
| DAQ Card | NI | PXI-6259 | 1 |
| DAQ Card | NI | PXI-6733 | 1 |
| Shielded Terminal Block | NI | SCB-68A | 3 |
| Multifunction Cable | NI | SH68F-68F-EPM | 1 |
| Multifunction Cable | NI | SHC68-68-A2 | 2 |
| Stage Controller | PI | C-884.6DC | 1 |
| Linear Stage | PI | M-406.4PD | 1 |
| XYZ Rot Stage | PI | M-900K276 | 1 |
| Optical Table | TMC | OPT.TOP/40”X96”x18” | 1 |
| Vibration Isolation | TMC | UltraDAMP | 1 |
| Computer | Colfax International | | 1 |
| Filter Wheel Controller | Sutter | LB10-3 | 1 |
| Vertical Gantry Post | Thorlabs | XT95-1000 | 2 |
| Horizontal Gantry Post | Thorlabs | XT95-500 | 1 |
| Rail Carrier | Thorlabs | XT95RC4 | 3 |
| Gantry Mount | Thorlabs | XT95P3 | 2 |
| Optical Table | TMC | 784-Special, 40”x96”x18” | 1 |
| Isolators | TMC | 14-424-39UD | 1 |
| **Macroscale Module** | | | |
| Galvanometer | Thorlabs | GVS012 | 1 |
| Electro-tunable Lens | Optotune | EL-16-40-TC-VIS-5D-1-C | 1 |
| Macro Zoom Body | Olympus | MVX-ZB10 | 1 |
| 1X Macro Objective | Olympus | MVXPLAPO 1X | 1 |
| Tube Lens Unit | Olympus | MVX-TLU-MVX10 | 1 |
| Camera | Hamamatsu | C13440-20CU | 1 |
| Emission Filter Wheel | Sutter | LB10-W32 | 1 |
| Excitation Objective | Nikon | 50 mm F1/4 G DLSR lens | 1 |
| Zoom Servo Motor | Dynamixel | MX-28R | 1 |
| **Nanoscale Module** | | | |
| Multi-Immersion Objectives | Applied Scientific Inc. | 54-12-8 | 2 |
| Remote Focusing Objective | Nikon | CFI S Plan Fluor LWD 20XC, NA 0.7 | 1 |

| | | | |
|---|---|---|---|
| Resonant Galvanometer | Cambridge Technology | CRS 4 KHz | 1 |
| Camera | Hamamatsu | C15440-20UP | 1 |
| Voice Coil | Thorlabs | BLINK | 1 |
| Half-Wave Plate | Newport | 10RP52-1B | 1 |
| Quarter Wave Plate | Bolder Vision Optik | AQWP3 | 1 |
| Polarizing Beam Splitter | Newport | 10FC16PB.7 | 1 |
| Emission Filter Wheel | Sutter | LB10-W32 | 1 |
| Stage Controller | PI | E-709 | 1 |
| Objective Piezo | PI | P-726.1CD | 1 |
| Breadboard | Thorlabs | B1824F | 1 |
| 95 mm Rails | Thorlabs | XT95SP-1000 | 2 |
| Rail Carriage | Thorlabs | XT95RC2 | 4 |
| Platform Positioner | Thorlabs | XT95N | 2 |

**Table S1** – Major hardware list for the Multiscale Cleared Tissue Axially Swept Light-Sheet Microscopy (MCT-ASLM). This table provides a detailed list of equipment used in the MCT-ASLM setup, including items shared between the macroscale and nanoscale modules (labeled as "common"). The table includes a description of each part, along with the manufacturer, product number, and quantity required. Note that this list is not comprehensive and serves as a general guide for the critical components used in the system.

| Device | Name | I/O Type | Purpose | Connector | Pinout (+/-) |
|---|---|---|---|---|---|
| **National Instruments PXI-6259** | | | | | |
| | P0.0 | DO | Macroscale Shutter | 0 | 52/18 |
| | P0.1 | DO | Master Trigger Out | 0 | 17/50 |
| | P2.0 | DO | Nanoscale Shutter | 0 | 37/4 |
| | CTR0 OUT | DO | Camera Trigger Out | 0 | 2/36 |
| | PFI0 | DI | Trigger Source | 0 | 11 |
| | AO0 | AO | Linear Galvo | 0 | 22/56 |
| | AO1 | AO | Resonant Galvo | 0 | 21/55 |
| | AO2 | AO | Electro-Tunable Lens | 1 | 22/56 |
| | AO3 | AO | Voice Coil | 1 | 21/55 |
| **National Instruments PXI-6733** | | | | | |
| | P0.0 | DO | Laser Port Switcher | 0 | 52/18 |
| | P0.2 | DO | 488nm Laser Enable | 0 | 49/15 |
| | P0.3 | DO | 561nm Laser Enable | 0 | 47/13 |
| | P0.4 | DO | 642nm Laser Enable | 0 | 19/53 |
| | AO0 | AO | 488nm Laser Intensity | 0 | 22/56 |
| | AO1 | AO | 561nm Laser Intensity | 0 | 21/55 |
| | AO2 | AO | 642nm Laser Intensity | 0 | 57/23 |
| | AO3 | AO | E-709 Digital Piezo Controller | 0 | 25/59 |
| **Computer Mainframe** | | | | | |
| | COM9 | Serial | Dynamixel Servo | | |
| | COM2 | Serial | Sutter Filter Wheel | | |
| | COM6 | Serial | Equipment Solutions | | |
| | | USB | C-884 Motor Controller | | |

**Table S2** – MCT-ASLM hardware connectivity. Analog and digital signals delivered from a National Instruments PXIe chassis equipped with PXI-6259 and PXI-6733 boards. Name, input/output (I/O) type, purpose, connector type, and pinout details are provided. The PXI-6259 board features two connectors, both of which are listed. Additional devices were configured to communicate via USB ports available on the computer mainframe.

| Magnification | Number of beads | Resolution (nm) | | |
|---|---|---|---|---|
| | | X | Y | Z |
| **Before Deconvolution** | | | | |
| 1x | 1599 | 1232 (w=0.71) | 1195 (w=0.76) | 5479 (w=0.61) |
| 2x | 10079 | 1397 (w=0.65) | 1364 (w=0.63) | 5246 (w=0.71) |
| 3x | 6651 | 1499 (w=0.64) | 1466 (w=0.55) | 4485 (w=0.73) |
| 4x | 4258 | 1786 (w=0.60) | 1796 (w=0.57) | 4207 (w=0.74) |
| 5x | 2670 | 1743 (w=0.62) | 1846 (w=0.98) | 3975 (w=0.77) |
| 6x | 1807 | 2265 (w=0.98) | 2469 (w=0.55) | 4243 (w=0.74) |
| 32x | 8478 | 477 (w=0.77) | 472 (w=0.82) | 466 (w=0.62) |
| **After Deconvolution** | | | | |
| 6x | 2043 | 1870 (w=1) | 2090 (w=1) | 3200 (w=1) |
| 32x | 6996 | 393 (w=1) | 366 (w=1) | 333 (w=1) |

**Table S3** - Resolution measurements for the MCT-ASLM in aqueous contexts ($\eta$=1.333) determined using sub-diffraction beads embedded in 1% agarose. The nanoscale path used 200 nm beads, while the macroscale path used 1000 nm beads. Beads were localized and fit to a 3D Gaussian distribution, with the resulting Full-Width Half-Maximum values binned into a histogram. The raw, undeconvolved data were analyzed using a Gaussian mixture model with two populations, whereas for the deconvolved data, a single Gaussian population was sufficient to describe the distribution. For the mixture model, only the smaller resolution population, which likely represents single, isolated beads, is reported. The value "w" indicates the weight of this population in the Gaussian mixture model. Note: Only the resolutions measured at 6X and 32X magnification are Nyquist sampled. The remainder of the values are intended for qualitative comparison only.

| Magnification | Number of aggregates | Resolution (nm) | | |
|---|---|---|---|---|
| | | X | Y | Z |
| **Before Deconvolution** | | | | |
| 6x | 11 | 1490 | 2170 | 5930 |
| 38x | 44 | 487 | 498 | 599 |

**Table S4** - Resolution measurements for the MCT-ASLM in high-refractive index contexts ($\eta$ = 1.56, BABB) were determined using CF555-labeled antibody aggregates in chemically cleared lung specimens. Aggregates were localized, and line profiles in the X, Y, and Z dimensions were fit to a Gaussian distribution to assess resolution. Measurements were performed using antibody aggregates due to the tendency of BABB to dissolve fluorescent nanospheres, and the resolution is expected to be ~11% worse owing to the red-shifted emission of CF555 aggregates compared to YG polystyrene carboxylate microspheres.

| Antibody (Gene) | Conjugate | Host | Catalog and RRID | Dilution | Manufacturer |
|---|---|---|---|---|---|
| Tyrosine Hydroxylase (TH) | | Rabbit | AB152 | 1:200 | Sigma-Aldrich |
| Tubulin $\beta$ 3/TUJ1 (TUBB3) | | Mouse IgG2a | 801202 RRID: AB_2313773 | 1:250 | BioLegend |
| CD31 (PECAM1) | | Rat | 550274 RRID: AB_393571 | 1:50 | BD Pharmingen |
| Tubulin $\beta$ 3/TUJ1 (TUBB3) | AlexaFluor 594 | Mouse IgG2a | 801208 RRID: AB_2650636 | 1:250 | BioLegend |
| Alpha-Smooth Muscle Actin (ACTA2) | AlexaFluor 488 | Mouse IgG2a | 53-9760-82 RRID: AB_2574461 | 1:250 | Thermo Fisher Scientific |
| Podocin (NPHS2) | | Rabbit | PA579757: RRID: AB_2746872 | 1:250 | Thermo Fisher Scientific |
| Synapsin 1 (SYN1) | | Rabbit | 51–5200 RRID: AB_2533909 | 1:250 | Thermo Fisher Scientific |
| Pan Cytokeratin (CK4, CK5, CK6, CK8, CK10, CK13, CK18) | | Mouse IgG1 | C2931-.2ML RRID: AB_258824 | 1:250 | Sigma-Aldrich |
| Six2 (SIX2) | | Rabbit | 11562-1-AP RRID: AB_2189084 | 1:250 | Proteintech |
| Endomucin (EMCN) | | Goat | AF4666 | 1:100 | Fisher Scientific |
| Laminin Subunit $\alpha$1 (LAMA1) | | Rabbit | L9393 RRID:AB_477163 | 1:125 | Sigma-Aldrich |
| Firefly Luciferase | | Goat | Ab181640 RRID:AB_2889835 | 1:250 | Abcam |
| Green Fluorescent Protein | | Chicken | AB_2307313 | 1:100 | Aves Labs |
| Anti-Chicken IgG (H+L) | AlexaFluor 647 | Donkey | 703-606-155 RRID: AB_2340380 | 1:300 | Jackson ImmunoResearch |
| Anti-Goat IgG (H+L) | AlexaFluor 555 | Donkey | A21432 RRID: AB_2535853 | 1:200 | Thermo Fisher Scientific |

| | | | | | |
|---|---|---|---|---|---|
| Anti-Goat IgG (H+L) Cross-Adsorbed | AlexaFluor 594 | Donkey | A-11058 RRID:AB_2534105 | 1:200 | Thermo Fisher Scientific |
| Anti-Rabbit IgG (H+L) Highly Cross-Adsorbed | AlexaFluor 647 | Donkey | A-31573 RRID:AB_2536183 | 1:200 | Thermo Fisher Scientific |
| Anti-Mouse IgG2a (H+L) Cross-Adsorbed | AlexaFluor 647 | Goat | A-21241 RRID:AB_2535810 | | |
| Anti-Mouse IgG2a (H+L) Cross-Adsorbed | AlexaFluor 488 | Goat | A-21131 RRID: AB_2535771 | | |
| Anti-Mouse IgG1 (H+L) Cross-Adsorbed | AlexaFluor 488 | Goat | A-21121 RRID:AB_2535764 | | |
| Anti-Rabbit IgG (H+L) Highly Cross-Adsorbed | AlexaFluor 568 | Donkey | A10042 RRID:AB_2534017 | | |
| Anti-Rat IgG (H+L) Highly Cross-Adsorbed | AlexaFluor 647 | Donkey | A78947 RRID:AB_2910635 | | |

**Table S5** –Details of primary and secondary antibodies used in this study. The table includes information on the antibody antigen (gene in parentheses), host species, catalog number, RRID (if available), working dilution, and manufacturer. For secondary antibodies, the fluorescence label is also specified.

| Reagent | Manufacturer | Product Number |
|---|---|---|
| 32% Paraformaldehyde (formaldehyde) aqueous solution | Electron Microscopy Sciences | 15714 |
| Dimethyl sulfoxide | Sigma-Aldrich | D8418 |
| Methanol | Fisher Scientific | A433-P |
| Dichloromethane | Sigma-Aldrich | 270997 |
| Dibenzyl Ether | Sigma-Aldrich | 33630 |
| Benzyl Alcohol | Sigma-Aldrich | 108006 |
| Benzyl Benzoate | ThermoFisher Scientific | 105860010 |
| Agarose | Fisher | BP160-500 |
| Donkey Serum | Jackson ImmunoResearch | 017-000-121 |
| Sheep Serum | Gibco | 16070096 |
| 10x Casein Solution | Vector Laboratories | SP-5020-250 |
| Igepal CA-630 | Sigma-Aldrich | I8896 |
| Triton X-100 | Fisher | BP151-500 |
| Nonidet P 40 | Sigma | 74385 |

**Table S6** – Chemical Information.

| Experimental Models | Source | Strain |
|---|---|---|
| Swiss Webster | Taconic Biosciences | Tac:SW |
| C57Bl6/J | The Jackson Laboratory | 000664 |
| Sprague Dawley | Charles River | 400 |
| NOD.Cg-Prkdc$^{scid}$ Il2rg$^{tm1Wjl}$/SzJ | The Jackson Laboratory | 005557 |
| Rosa26 PF4-Cre[41] | | Rosa26-CAG-loxp-stop-loxp-EGFP |

**Table S7** – Model Organism Information.
